# Reducing Movement Synchronization to Increase Interest Improves Interpersonal Liking

**DOI:** 10.1101/2021.06.30.450608

**Authors:** Inbal Ravreby, Yoel Shilat, Yaara Yeshurun

## Abstract

Synchronization has been identified as a key aspect of achieving mutual goals and enhancing social bonding. While synchronization could be maximized by increasing the predictability of an interaction, such predictability is in tension with individuals’ level of interest, which is tied to the interaction’s complexity and novelty. In this study, we tested the interplay between synchronization and interest. We asked 104 female dyads to play the Mirror Game, in which they were instructed to move their hands as coordinately as possible, and then report how much they liked each other. Utilizing information theory and video processing tools, we found that a combination of movement synchronization and complexity explained liking almost two times better than movement synchronization alone. Moreover, we found that people initiated novel and challenging interactions, even though they paid a price – being less synchronized. Examining the interactions’ dynamics, we found that people who liked each other moved in a more synchronized, complex, and novel manner during most of the interaction. This suggests that in addition to synchronization, maintaining interest may be critical for positive social bonding. Thus, we propose a new framework in which balancing synchronization and interest, rather than merely maximizing synchronization, optimizes the interaction quality.

## Introduction

Sensorimotor communication, which is achieved through modification of individuals’ body movements to communicate their intentions, is integral to human social communication and coordination (1,2) as well as in various other animals (3–6). It has been shown that during social interactions, dyadic coordination and synchronization of movements play a key role in successful interaction and achievement of mutual goals (7,8). As sensorimotor communication is a multifaceted process, in this study we explored aspects beyond synchronization that lead to positive social interactions in humans following a coordination task – movement complexity and novelty.

### Synchronization Increases Social Interaction Quality and Vice Versa

A large body of literature has shown a positive relationship between the quality of social interaction and synchronization of movements (9–11). Synchronization of movements is defined as the similarity of movements (12), either in the time domain or the frequency domain (11). For example, when individuals interacted with others who mimicked their body language, synchronization of movements was shown to increase liking (9), affiliation (13), rapport (14), trust (11,15) and collaboration (16). These results are related to a broader concept of synchrony that was offered as a mechanism for social understanding (17,18). Specifically, it was suggested that movement synchronization is a body-based way for creating socioemotional connection and shared experiences (18,19). In coordinated actions, or when mirroring each other, the actions of self and the other overlap, and this facilitates mutual understanding (20). Synchronization of body movements is also related to positive feelings such as connectedness and togetherness (13,19,21,22). Remarkably, there is not only a correlation, but also a bidirectional causality between the social interaction quality and synchronization (21,23,24). The aforementioned studies have established the role movement synchronization plays in successful social interactions.

### Predictable Movements are Easier to Sync with, but are not Interesting

Making oneself predictable may be used as a synchronization strategy. For example, when two acquaintances are approaching one another and want to greet each other using a physical gesture, one person may intend a handshake, while the other may intend a hug. If one of the acquaintances starts raising the right hand when approaching the other, the partner would know in advance to match their movements so they will shake hands successfully. Indeed, research shows that predictable movements facilitate human synchronization during non-verbal interaction (25–27). Thus, if individuals make predictable movements, their coordination increases since they can plan how to move properly, and coordinate their movements. Increasing predictability, therefore, may increase the likelihood of positive social interaction.

Notably, a predictable movement does not have to be repetitive, and vice versa. A movement may be new but simple, or it may be complex but repetitive. Significantly, a highly predictable or repetitive interaction is not necessarily rewarding, as the interaction might feel tedious (28,29) and boring (30). Boredom may be defined as the aversive experience of wanting, but being unable to engage in a satisfying and challenging activity (28,31). Any dyadic interaction can be conceived as a dynamic process by which information is exchanged between individuals (32). Following that, boredom may occur when there is a lack of rich information, making the activity unsatisfying. Such an experience is likely to occur during highly predictable and repetitive interactions. One can avoid boredom by exploring novel stimuli – that is, a change in stimuli conditions from previous experience (33). Moreover, individuals constantly strive for complexity and novelty (29,34), as such patterns violate prior expectations, and thus may lead to more interesting and challenging interactions. The more patterns are unpredictable (complex) and less repetitive (novel), the more rewarding the interactions. This notion is consistent with Hasson and Frith (32) who argued that we need to go beyond simple mirror alignment once we start interacting, and with Wohltjen and Wheatley who demonstrated the intricate dynamics of pupil dilation synchronization (35).

### The Current Study

Altogether, we suggest a novel framework in which in order to optimize the interaction quality, there should be a subtle balance between being synchronized and generating an interesting interaction. This implies that in contrast to the current view, maximizing the synchronization per se would not necessarily result in better interaction. Accordingly, in this study we set out to test the role of synchronization, complexity, and novelty in generating positive interactions, in the context of sensorimotor dyadic communication. We hypothesized that interesting movements, which are complex and novel, will improve the interaction more than mere synchronization. Moreover, we hypothesized that when people are asked to synchronize their movements with each other, they will choose to add complexity and novelty to the interaction, despite greater difficulty in arriving at synchronization.

## Methods

### Participants

Twenty-six naïve females participated in this study. The participants were from Rehovot, Israel, or nearby. We recruited the participants using social media. To validate that they do not know each other we conducted preliminary phone interviews. We recruited only females since the link between movement synchrony and affect has been found to be strongest in female dyads (36). Among the participants in this study, 10 have originally participated in another study that is yet to be published. Importantly, its analyses are not related to this study. The experiment took place over three different sessions: the first session included six participants (mean age = 26.83, *SD* = 4.21 years); the second session included ten participants (mean age = 25.6, *SD* = 3.1 years); the third session included ten participants (mean age = 26, *SD* = 3.46 years). In each round participants were randomly assigned partners and played the Mirror Game (see details below) for about two minutes in a round-robin design, i.e., each participant played the game with all of the other participants in the same session. Thus, there were 15 dyads in the first session, 45 dyads in the second session and also 45 dyads in the third session. The duration of the Mirror Game ranged between 109.81 to 130.56 seconds (*mean* = 119.84, *SD* = 2.5). All participants provided written informed consent and were paid for their time. We excluded one dyad from the analyses, due to lack of movement during the game. Following that, a total of 104 dyads were included in the motion analyses. The sample size is in accordance with previous studies about the relationship between movement synchronization and rapport (e.g.,(14). According to previous findings the strength of the correlation between movement synchronization and liking is approximately equal to *r = .4* in a simple tapping task (in experiment 1, *r* = .39 and in experiment 2, *r* = .40; (13). However, because of the complexity of the Mirror Game task relative to a tapping task, we expected lower correlation coefficient ranging between r =0.25 and r=0.3. We ran power analysis for this range of expected effect sizes, which corresponds to a sample size of 85-123 dyads.

### Experimental Design

Participants were assigned to pairs and played the full-body Mirror Game (37), in which they were instructed to move their hands with as much coordination as possible for about two minutes, while keeping their legs in place, with no designated leader or follower (see Fig. 1a). Each dyad played the game in a separate room. During the game, participants stood at a distance of 50 cm apart, which was marked on the floor. Participants were not allowed to speak with each other during the entire experiment; thus only non-verbal factors influenced the impression formation. After each dyadic interaction, both players indicated on a visual analog scale ranging from 0 to 100 how much they liked their partner. These subjective reports were done via computers in a class with distributed computers, so no one could see the others' screen. Throughout each game, participants were filmed by a hidden camera alarm clock, which filmed them in 29.97 frames per seconds (FPS) and was set on a table in each of the five rooms used for the experiment, at a distance of 159 cm from where the players stood. After we completed the data collection, we used the video recordings to verify that all participants followed the instructions and moved during the whole game, mirroring each other without talking. In addition, we used the video recordings to analyze participants’ movement.

**Fig. 1.**
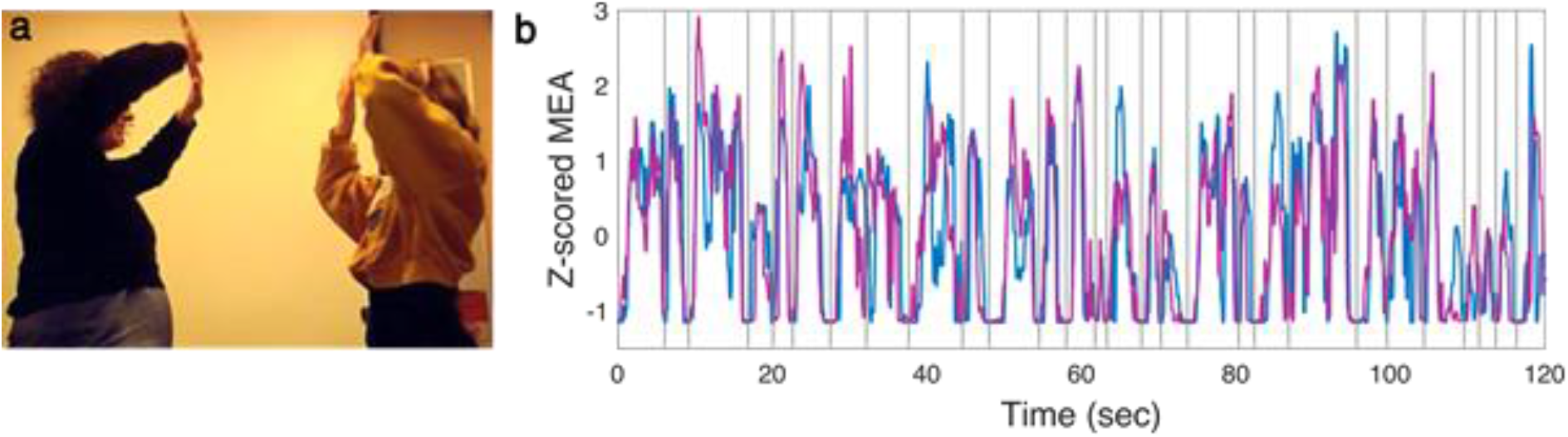
The Mirror Game. (a) A pair playing the Mirror Game while a hidden camera alarm clock, that was placed on a table next to the players, is filming the game. (b) The z-scored MEA signals of two representative participants while playing with each other. The vertical lines denote the separation between the movement segments.

### Motion Energy Analysis

We used Motion energy analysis (MEA) to quantify each player’s motion throughout the Mirror Game (38). Motion energy was defined as frame-by-frame differences in pixels color between consecutive video-frames (23,39). Since, throughout the recordings, both camera position and lighting conditions were kept constant, any frame-by-frame changes indicated body motion of the respective player and not of its surroundings. Following Ramseyer and Tschacher (23), we performed video-noise reduction using automatic detectors for time-series of raw pixel-change. The MEA signal of each player was transformed to z-scores to scale the final values and thus account for differences in the players' height and hand size (Fig. 2). We performed the analyses both at the level of the whole game and along movement segments (22).

**Fig. 2.**
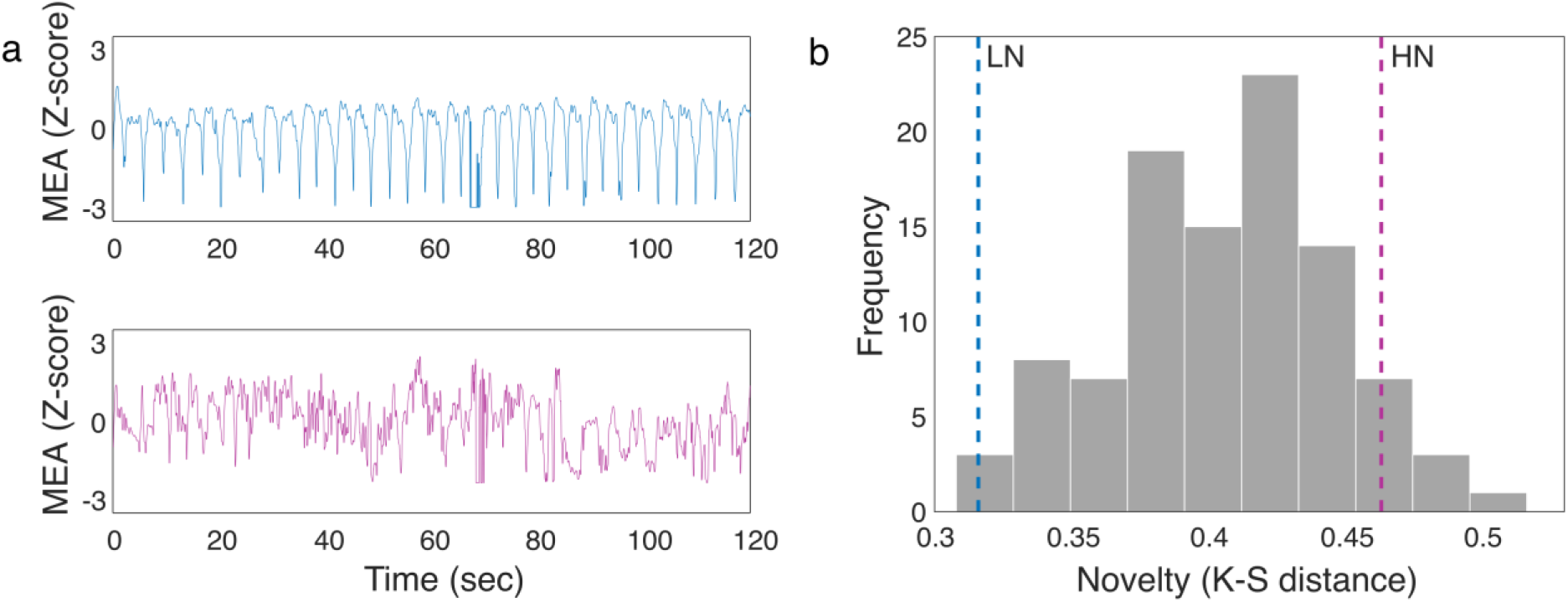
Moving in a novel manner. (a) The upper panel shows the z-scored MEA of repetitive up and down hand movements (LN). The bottom panel shows the z-scored MEA of diverse and novel hands movements (HN). The hand movements were measured in a setup identical to that of the dyadic Mirror Games. (b) A histogram of the novelty of the dyads’ movements during the Mirror Game. The blue dashed line denotes repetitive hand movements up and down (LN). The magenta dashed line denotes diverse movements (HN).

### Segmentation Into Movement Windows

In order to divide the z-scored MEA vectors of each dyad into movement segments, the signals were smoothed using moving average to eliminate short-term trends and to improve signal to noise ratio. The optimal window for the moving average was heuristically evaluated. Next, we searched for local minima within the signals, i.e., stopping and deceleration points, each having to meet the following conditions: (1) The value was smaller than zero (the average movement velocity); (2) Minimum points must be separated by at least the length of the optimal time window; (3) Two consecutive minimum points must differ by more than 0.1 SDs. After finding the minimum points in the signal of each participant in each dyad, we searched for the minimum points with the smallest time differences between the participants in each dyad and defined them as shared minimum points. Two consecutive minimum points had to be separated by at least the length of the optimal moving average window. To set a starting point and an end point for each movement, we averaged the time indices of minimum points in each dyad’s vectors (See Fig. 1b for a representative movement segmentation). The time window analysis was smoothed using a moving average of four seconds, which was the average segment length, to improve signal to noise ratio.

### Measuring Movement Synchronization

In accordance with previous studies, we used Pearson correlation to measure the synchronization level of dyads **(21)**, and thus measured the degree of accelerating or decelerating at the same time.

### Measuring Movement Complexity

We used Shannon entropy (40) – the average level of information – as a measure of the complexity of dyad’s movements. Although movement is a continuous signal, we obtained a sample every 0.033 seconds, so our data were discrete. The entropy of a discrete variable *X* that can take values in the set *A* is defined as:

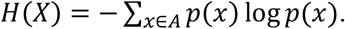

To calculate the entropy of each player’s z-scored MEA signal, we first calculated the frequency of each unique MEA value per sample, and then calculated the entropy using the Entropy function in MATLAB 2020a. Next, we averaged the entropies of the two participants in each dyad. This was done in order to get a single measure of dyadic entropy, which represents the dyadic average level of movement information. Communication is said to occur when information flow from one location to another and cause a change in the receiver (40). Information theory, the theoretical framework underpinning entropy, attributes higher levels of information to more unpredictable distributions. In other words, when an interaction is less predictable it is more complex (25,41). Therefore, unpredictable interactions are more informative and complex than predictable ones, and hence also more interesting.

### Measuring Movement Novelty

In addition to complexity, we developed a new method for quantifying dyadic novelty – how different (not-repetitive) each movement segment was compared to the previous segments. Using the Kolmogorov–Smirnov distance (K-S distance) test (42), we quantified the distance between the empirical distribution functions of the z-scored MEA of each segment to each of the previous segments. Each test consists of two segments at a time. Notice that the two-sample Kolmogorov–Smirnov test, which is used to test whether two samples come from the same distribution, is highly appropriate in our case, as it is sensitive to differences in both location and shape of the empirical cumulative distribution functions (CDFs) of each two segments. In general, the distance between empirical CDFs is small when the distributions are similar (it is zero when they are identical) and close to one when they are very different. We compared the first and the second segment. This was done for each player in each dyad. We then averaged the statists of the two players within each dyad to get a single dyadic statist. This averaged statist served as the novelty score of the second segment. Since there are no segments before the first one, it was also the novelty score of the first segment. Next, we compared the third segment and the second segment, and the third segment and the first segment. The average statist of the two comparisons, averaged also between the two players within each dyad, served as the novelty score of the third segment. Similarly, we compared the fourth segment and each of the previous segments, and averaged the three resulting statists of each player within each dyad, and also averaged between the two players to get the novelty score for the fourth segment. This was done for each dyad up to segment N. Thus, each segment was assigned a novelty score, which enabled tracking the novelty changes along the time domain. In order to have also a single novelty score for each dyad, we averaged the novelty scores of all the segments (without the first one, which got the same value as the second segment) within each dyad. We then compared it to a novelty scale we developed by filming a volunteer (who was not a part of the Mirror Game cohort) twice: once while periodically moving their hands up and down for two minutes in a repetitive manner, and once while moving their hands in the most innovative way possible, without repeating any movement patterns, for two minutes. This was done using the exact same setup used in the Mirror Game experiment. We defined these movements as low novelty movements (LN, upper panel of Fig. 2a) and high novelty movements (HN, bottom panel of Fig. 2a), respectively. We calculated the z-scored MEA of these two videos and then calculated their novelty (using the K-S distance).

In addition to objectively measuring the novelty of the movements, a judge watched each video of the Mirror Game, and was requested to decide whether the players had moved differently than simply moving their hands up and down repeatedly throughout the whole interaction (or otherwise in a highly repetitive and thus predictive manner, e.g. from side to side or outwards and then inwards).

### Measuring Liking

To estimate participants’ mutual liking, we calculated a dyadic score of likeability per each Mirror Game, which was defined as the average reported liking of the two players. We wanted to test whether movement synchronization, complexity, and novelty predict how much participants liked each other. To do so, we used multiple linear regression models and calculated the explained variance of liking by synchronization solely, synchronization together with complexity, and synchronization together with complexity and novelty. A total of three models were tested using JASP version 0.14.1.0 software. In addition, we calculated the Pearson correlation between synchronization, complexity, novelty and mutual liking. We excluded four dyads from the analyses, who were farther than 3 SDs from the average of each measure (two were outliers due to their complexity levels, and two due to their liking ratings). In total, the analyses were performed on 100 dyads.

### Granger causality analysis (GCA)

GCA is a method that uses autoregressive models to measure the causal relationship between two time series (43). We computed the pairwise-conditional causalities of each participant’s movement in the dyad to that of the other participant.

### Movement dynamics

To test whether dyads’ movement dynamics during the interaction reflected mutual liking, we divided the dyads according to the median into two groups: high-liking half and low-liking half, each comprising 50 dyads, and compared the dynamics of synchronization, complexity, and novelty between the two groups.

### Bayesian Analysis

We have provided a complementary measure to the standard *P* values (44), using Bayes factors (*BF*) calculated with the JASP software; priors were set according to default JASP priors (45). The interpretation of the Bayes factors was according to the following scale: Bayes factors between 3-10 provide substantial evidence against H_0_; Bayes factors between 10-100 provide strong evidence against H0; and Bayes factors above 100 provide decisive evidence against H_0_ (46). The Bayes factors calculated for the predictors of the regression models (*β*s’ weights) were inclusion Bayes factors, denoting the change in the prior to posterior probability inclusion odds when including the predictor (47)

## Results

### Dyads’ Movement Reflected Reciprocal Relationships

Remarkably, the GCA showed that in 95 out of the 100 dyads, each player’s movement was significantly causally (Granger caused) determined by the other player’s movement, i.e., there was a significant mutual Granger cause. These results suggest that the vast majority of the participants did not consistently play the role of leader or follower, but rather changed roles during the game (48). Such a symmetric and reciprocal relationship may be more rewarding (49), but may make it harder to be in sync with each other, as movement planning is harder, because the interaction is less predictable. The remaining five dyads exhibited a leader-follower movement pattern, as was shown by the one-way relationship of the Granger causality – in these five dyads one participant Granger-caused the other participant’s movement, and there was no reciprocity. Combining these two findings shows that in all the dyads at least one participant Granger-caused the movements of the other, and, therefore, the partner mirrored their movements, suggesting that all dyads followed the instructions and moved in a coordinated manner.

### Dyads Chose to be Novel and Complex

Our results indicate that among all 100 dyads, there was not a single dyad in which participants repeated the same hands movements throughout the whole Mirror Game, according to a judge who watched the videos of the Mirror Game. Moreover, the novelty score we developed (see methods for more details) revealed that 99% of the dyads moved in a more novel way than just moving the hands up and down repeatedly (LN), even though we had not instructed them to perform novel movements. Furthermore, only 9% of the dyads moved in a more novel manner than when intentionally moving the hands in a highly diverse manner (HN). This result further validates the novelty measure we chose, since our defined low-novelty measure (LN) was at the far-left side of the novelty histogram (.32, as marked by the blue dashed line in Fig. 2b) and our definition for high-novelty movements (HN) was at the right side of the novelty histogram (.46, as marked by the magenta dashed line in Fig. 2b). The mean novelty of all dyads = .4, *SD =* 0.04, and two-tailed one-sample t-test showed that low-novelty movements, repeated hands movements, were significantly lower than the average novelty *t*(99) = 21.84, *P* < .001, Cohen’s *d* = 2.18, 95% *CI* = [1.82, 2.54], BF_10_ > 1,000, whereas high-novelty movements, diverse hands movements, were significantly higher than the average novelty *t*(99) = 14.05, *P* < .001, Cohen’s *d* = 1.4, 95% *CI* = [1.68, 1.13], *BF*_*10*_ > 1,000. Thus, even though moving hands up and down repeatedly makes it easier to get into sync (26,27) and is in accordance with the instructions, 99% of the dyads preferred moving in a less repetitive, more novel, and more interesting manner.

Next, we tested the relationships between the synchronization, complexity, and novelty of the dyads. Synchronization was measured using Pearson correlation; complexity was calculated using Shannon entropy (40), and novelty was measured using K-S distance (see Methods section for more details). We found a significant negative correlation between the synchronization and complexity levels of the dyads, *r* = −.27, *P* = .007, 95% *CI* = [0.08, .44], *BF*_*10*_ = 4.252, and between their synchronization and novelty levels, *r* = −.32, *P* = .001, 95% *CI* = [−.48, −.13], *BF*_*10*_ = 21.88 (Fig. 4a-b). This supports the notion that in more complex and novel interactions, it is harder to get into sync. These results suggest that humans do not only seek to be in sync with others, even when they are instructed to do so, but rather they seek to avoid predictable and repetitive interactions. They strive to be synchronized in an interesting, complex, and novel manner. Notably, novelty was found to be significantly positively correlated with complexity, *r* = .39, *P* > .001, 95% *CI* = [.21, .54], *BF*_*10*_ = 344.21 (Fig. 4c).

### Synchronization and Complexity Play a Role in Mutual Liking

We used multiple linear regression models as well as Bayesian regression models to predict dyads’ mutual liking using synchronization, complexity, and novelty (see Methods section for more details). The mutual liking of dyads was calculated as the average reported liking of the two players (see Methods section for more details). A model that included only the synchronization level predicted 7.8% of variance in liking, *β* = .28, *t*(98) = 2.88, *P* = .005, *R*^2^ = .08, *BF*_*10*_ = 7.672 (Fig. 4a). This result is in line with previous studies (9), showing the linkage between synchronization and liking. Including complexity level into the model significantly improved it, explaining almost twice the variance of the former model. Accordingly, the linear model with both synchronization and complexity as predictors, predicted 14.9% of the variance in liking, *F*(2, 97) = 8.49, *P* < .001, *BF*_*10*_ = 66.54, with a positive correlation between synchronization level and liking, *β* = .35, *t*(97) = 3.63, *P* < .001, *BF*_*inclusion*_ = 47.973, and between complexity and liking, *β* = .28, *t*(97) = 2.85, *P* = .005, *BF*_*inclusion*_ = 13.857 (Fig. 4b). Notably, adding complexity level to the model increased the predictive power of synchronization, specifically in its Beta weight. This indicates that adding complexity to the model suppressed the irrelevant variability of synchronization and thus further improved the model. Introducing novelty to the model did not significantly improve it, and predicted an additional variance of 0.05%, *P* = .456, *BF*_*10*_ = .37. This result suggests that although the average novelty during the game is related to the synchronization level and to the complexity of the interaction, it is not sufficient to create affiliation towards others. Following the comparisons between the models, as demonstrated in Fig. 4b (and in Supplementary Material Table 1), the best linear model out of the three was:

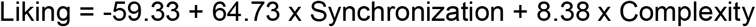

Similar results were also found using maximal cross-correlation as the synchronization measure (see Supplementary Material Fig. 1).

Moreover, we conducted a stepwise multiple linear regression on all available independent measures (see Supplementary Material for further details) to evaluate the effect of the order of the variables that were entered to the regression model. This analysis yielded almost identical results, therefore we are presenting here the results of the multiple linear regression, which is a more prominent method of analysis. Taken together, there is a converging evidence that individuals tend to like others when their interactions are more complex and unpredictable, rather than simple and highly predictable.

### Dynamics of Synchronization, Complexity and Novelty in Dyads with High and Low Mutual Liking

We were interested in testing the relationship between the dynamics of dyads’ movements during the interaction and liking. To test this, we compared the dynamics of synchronization, complexity, and novelty between the high-liking and low-liking groups (see Fig. 5), both throughout the entire game and along movement segments (22) (see Methods section for more details). As for synchronization (Fig. 5a), in 55.87% of the total game, the high-liking group moved in a more synchronized manner than the low-liking group. Binomial sign test shows that this result was significantly above chance level (i.e. 50%), binomial *P* < .001. Moreover, the difference between the groups was significant (*P* < .05) for 1.2% of the game duration, such that the high-liking group was less synchronized than the low-liking group. Two-tailed Bayesian analysis showed no substantial evidence (*BF*_*10*_ < 3) that the high-liking group is less synchronized than the low-liking group. Similar results were also found using maximal cross-correlation as the synchronization measure (see Supplementary Material Fig. 2).

Regarding complexity (Fig. 5b), during 79.62% of the game, the movements of the high-liking group were more complex than those of the low-liking group, which is significantly above chance level, as shown by a binomial sign test, *P* <.001. Furthermore, the difference between the high and low-liking groups was significant for 15.66% throughout the duration of the game – the high-liking group moved in a more complex manner than the low-liking group. Finally, this was substantially supported by the Bayesian analysis in 4.01% of the Mirror Game duration.

In terms of novelty (Fig. 5c), unlike in the results of the multiple linear regression model, which did not take into account the time domain, in 90.16% of the game the high-liking group moved in a more novel manner than the low-liking group, binomial sign test *P* < 0.001. Moreover, the difference between the high- and low-liking groups was significant for 10.7% of the game – the high-liking group was more novel than the low-liking group. Using Bayesian analysis, this was substantially supported in 1.94% of the duration of the Mirror Game.

As shown in Fig. 5c, it seems that novelty increased over time, both in the low and the high-liking groups, but more prominently in the low-liking group. To test this, we performed a post-hoc correlation analysis and found that, indeed, the novelty significantly increased over time in the low-liking group, *r* = .82, *P* = .001, 95% *CI* = [.70, .89], *BF*_*10*_ > 1,000, and in the high-liking group, *r* = .67, *P* =.001, 95% *CI* = [.48, .80], *BF*_*10*_ > 1,000. Remarkably, a comparison between these Pearson coefficients showed that the increase in novelty over time of the low-liking group was significantly larger than that of the high-liking group, *z(96)* = 1.69, *P* = .045. This was also demonstrated by the slopes of the fitted linear regression lines: Novelty_low-liking_ = 6.34×10^−4^ × Time (s) + .35, Novelty_high-liking_ = 3.29×10^−4^ x Time (s) + .39. Notably, although the novelty of the high-liking group increased more slowly over time, the level of novelty in this group was relatively high right from the beginning of the Mirror Game, suggesting that people who like each other tend to create novel interactions from the very beginning, and then gradually increase the novelty level.

## Discussion

In this study we demonstrated that in addition to synchronization, complexity and novelty play a role in generating positive interactions. We also found that when they were synchronized, participants who performed more complex and novel movements while playing the Mirror Game, liked each other more. Our results suggest that humans seek complexity and novelty, even when getting into sync will be more difficult.

In line with previous studies demonstrating that movement synchronization between individuals increases the positivity of social interactions (9,11), we found that elevated synchronization during the Mirror Game was associated with elevated mutual liking (Fig. 4, 5a). Interpersonal synchrony may signal social proximity or similarity (17) and plays an important role in social cohesion (18,50). It was suggested that in small groups such as dyads, greater attention to the movements of each other, and synchronized movements, or mirroring each other, lead to a blurring between the self and the other (13,51). In turn, this increases the feeling of ‘being in the zone’ which is also known as togetherness (13,19,21,22). Moreover, it was suggested that being synchronized is rewarding because it is an effective way to understand one’s interaction partner (17). Consequently, such an alignment may facilitate communication (52).

When playing the Mirror Game, the only instruction the dyads received was “to move their hands with as much coordination as possible, mirroring each other”. It is easier to be in sync when the movements are more predictable (25–27) and repetitive, and when there is a leader-follower type of interaction, with fixed roles. Although it made it harder for the participants to meet the single requirement of the experiment, in almost all of the Mirror Games (95%), both players influenced each other’s motion, as was demonstrated by the GCA results. This occurs when two participants reciprocally adjust their ongoing rhythms as a result of an interaction, serving as a reliable marker for mutual sharing of information (53,54). This is in contrast to the leader-follower type of interaction, which contains only a one directional information flow. In addition, the dyads chose to perform complex and novel movements, rather than merely performing synchronized movements. Accordingly, the results suggest that participants preferred to decrease their potential synchronization, in order to increase the mutual interest and experience a better, more reciprocal interaction. This may imply that strong relationships between mutual influence and positive interactions (55) and between elevated interest and positive interaction (56) are fundamental for human beings.

Taking into account the whole interaction, we found that the synchronization and complexity of movements predicted liking more accurately than only synchronization of movements (Fig. 4). Adding complexity to the model suppressed the irrelevant variability of synchronization and further improved the prediction of liking. In addition, synchronized and complex movements were negatively correlated (Fig. 3a). This finding implies that liking relies on a tradeoff between the synchronization and the complexity of an interaction. In other words, the results support the notion that on one hand people seek predictable interaction which facilitates synchronization, and in turn is rewarding (17), but on the other hand, people seek interesting and challenging interactions. Complex interactions are by definition less predictable and comprise a high level of information transformation during communication (as mentioned regarding the definition of entropy). Thus, an optimal balance between the two may maximize the reward and the quality of the interaction.

**Fig. 3.**
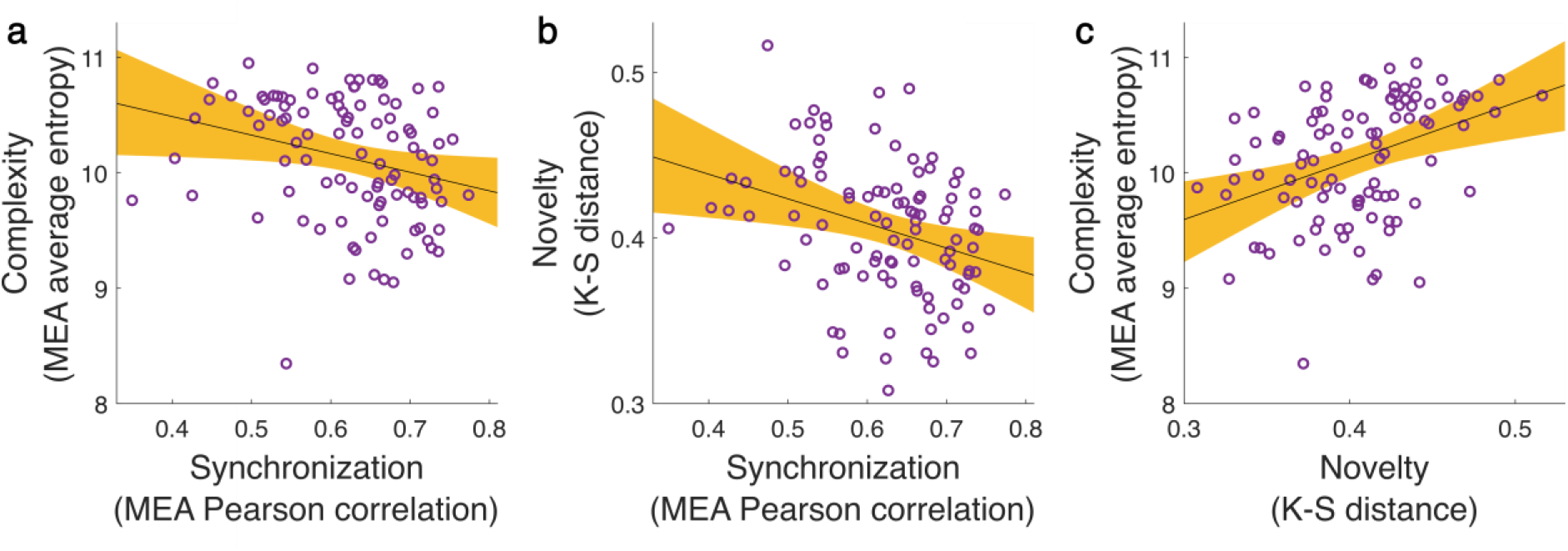
A tradeoff between synchronization and being interested. Pearson correlations between the different movement measures. In each panel each circle represents a dyad, and the black line is the linear regression line. The orange area marks the confidence interval around the slope of a regression line. (a) Negative correlation between complexity and synchronization, *r* = −.27, *P* = .007, 95% *CI* = [−.44, −.08], *BF*_*10*_ = 4.252. (b) Negative correlation between novelty and synchronization, *r* = −.32, *P* = .001, 95% *CI* = [−.48, −.13], *BF*_*10*_ = 21.88. (c) Positive correlation between complexity and novelty, *r* = .39, *P* < .001, 95% *CI* = [.23, .54], *BF*_*10*_ = 344.21.

**Fig. 4.**
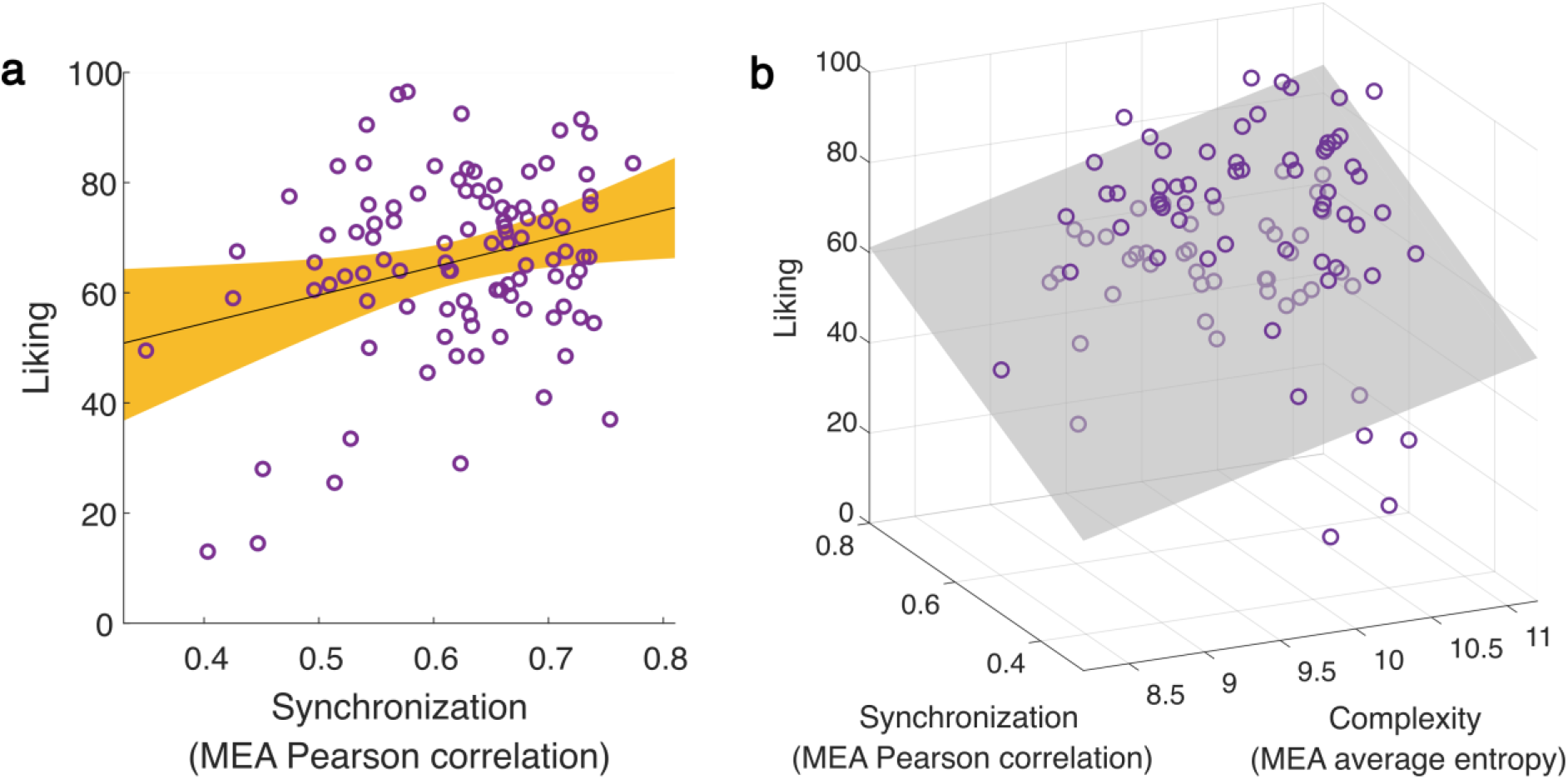
Predicting liking by dyadic movement features. (a) Synchronization level of the z-scored MEA signals significantly predict liking, *β* = .28, *t*(98) = 2.88, *P* = .005, *R*^2^ = .08, BF_10_ = 7.672. Each circle represents a dyad, the black line is the linear regression line, and the orange area marks the confidence interval around the slope of a regression line. (b) Multiple linear regression model including both synchronization and complexity significantly improved the model predictions, this model predicted 14.9% of the variance in liking, *R*^2^ = .149, *F*(2, 97) = 8.49, *P* < .001, *BF*_*10*_ = 66.541. Each circle represents a dyad in the 3D space and the grey plane marks the regression surface that was fitted by the model.

In contrast to the complexity of movements, when adding the average novelty of the interaction to the model, there was no improvement in predicting liking. Possibly, this is because novelty seeking is a fundamental drive (29,33,34) and, accordingly, may be a basic need in social interaction. Subsequently, it is plausible that during the Mirror Game as a whole, the dyads played in a novel manner, regardless of the level of liking. In 99% of the cases, the movements of the dyads were more novel than just repetitively moving their hands up and down, even though there was no requirement to move in a novel manner, which makes it harder to be in sync. Moreover, novelty increased over time, and this trend was even more pronounced in the low-liking group than in the high-liking group, which was relatively novel from the beginning of the game.

Our results showed that the dynamics of synchronization along the Mirror Game segments, unlike the synchronization of the dyad’s movement across the entire game, was only moderately related to liking (Fig. 5a). This discrepancy between the game as a whole and the dynamics on a movement-to-movement basis suggests that, in this case, the whole is greater than its parts, and to identify differences in synchronization, one has to consider the interaction as a whole. During the Mirror Game, people got in and out of sync, in accordance with previous findings (57). There were fluctuations in the level of synchronization along the movement segments, but, on the whole, people who liked each other were more synchronized. As Hasson and Frith (32) claimed, interacting individuals are dynamically coupled rather than simply aligned. Interactions are dynamic states which involve continuous mutual adaptation during complementary behavior develops. This notion was recently demonstrated in the context of pupil dilation synchronization during a conversation, which was found to fluctuate according to the shared attention pattern (35). Such continuous mutual adaptation along the time domain generates synchrony and, in doing so, promotes shared understanding (58). We suggest that in order to increase the interaction complexity and novelty, humans are willing to sacrifice synchronization to some extent, as a part of mutual adaptation. Together, complexity and novelty make an interaction interesting and meaningful – an interaction that one would want, unlike a highly synchronized, but very predictable, repetitive, boring interaction. This notion was supported by the positive relationship between the complexity and novelty of movements along the time domain, to liking. We suggest that when people feel confident enough to allow themselves to potentially risk losing synchronization, they try to increase the level of complexity and novelty to make the interaction even more satisfying. Then, they restore the synchronization and subsequently balance the two back and forth.

**Fig. 5.**
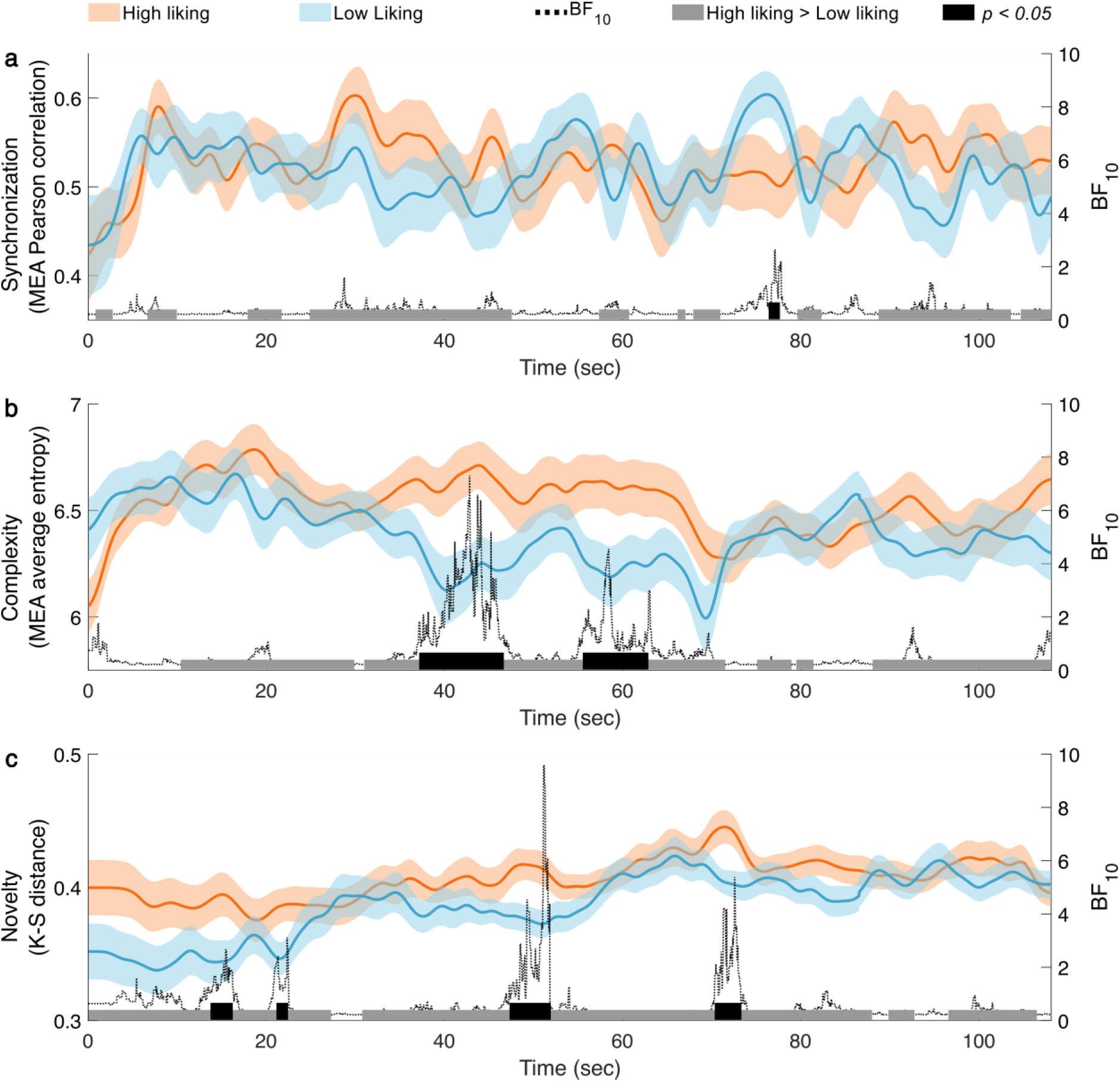
The dyadic movement measures along the interaction segments for high and low-liking. The orange and blue lines denote the average value of the high and low-liking groups respectively, along the Mirror Game time-points. The shaded orange and blue marks denote the *SE*s of the high and low-liking groups, respectively. The gray marks at the bottom of each panel denote time-points in which the high-liking group had a higher value than the low-liking group. The black marks denote time-points in which there was a significant difference between the liking groups. The dashed lines depict two-tailed Bayes factors (*BF*_*10*_). (a) Synchronization was higher in the high-liking group than the low-liking group for 55.87% of the game duration, Binomial sign test *P* < .001. (b) Complexity was higher in the high-liking group than in the low-liking group for 79.62% out of the game duration, Binomial sign test *P* < .001. (c) Novelty was higher in the high-liking group during 90.16% of the game, binomial sign test *P* < .001.

## Conclusion

To summarize, in addition to the canonical view that the interaction quality relies on the link between synchronization and prosocial emotions (21,59), we argue that complexity and novelty are crucial and even indispensable factors in human social interactions. Accordingly, we propose a new framework in which optimizing the interaction quality requires a delicate balance between being synchronized and generating challenging and interesting interaction.

## Acknowledgements

We would like to thank Yuval Hart for fruitful discussion. Y.S would like to thank Prof. Avishai Henik and Dr. Moti Salti for their support during this project. I.R. was supported by a fellowship from the Ariane de Rothschild Women’s Doctoral Program. This research was supported by the Israel Science Foundation (grant No. 2434/19).

## Author Contributions

Conceived the idea: IR; YS. Designed experiments: IR. Ran experiments: IR. Analyzed data: IR; YS. Wrote first draft: IR, YS. Edited final draft: IR; YS; YY. Supervision: YY.

## Data availability

The code is available from the corresponding authors upon request. The data that support the findings in this study are available at https://osf.io/6pbgf/?view_only=b02268caee0a4ee4ae728d14bcbb5a65.

## References

1. Ansuini C, Cavallo A, Bertone C, Becchio C. The visible face of intention: why kinematics matters. Front Psychol [Internet]. 2014 [cited 2020 Jun 13];5. Available from: https://www.frontiersin.org/articles/10.3389/fpsyg.2014.00815/full

2. Pezzulo G, Donnarumma F, Dindo H. Human Sensorimotor Communication: A Theory of Signaling in Online Social Interactions. Daunizeau J, editor. PLoS ONE. 2013 Nov 20;8(11):e79876.

3. Rands SA, Cowlishaw G, Pettifor RA, Rowcliffe JM, Johnstone RA. Spontaneous emergence of leaders and followers in foraging pairs. Nature. 2003 May;423(6938):432–4.

4. Sirot E, Touzalin F. Coordination and Synchronization of Vigilance in Groups of Prey: The Role of Collective Detection and Predators’ Preference for Stragglers. The American Naturalist. 2009 Jan 1;173(1):47–59.

5. Cote J, Bocedi G, Debeffe L, Chudzińska ME, Weigang HC, Dytham C, et al. Behavioural synchronization of large-scale animal movements – disperse alone, but migrate together? Biological Reviews. 2017;92(3):1275–96.

6. Bode NWF, Faria JJ, Franks DW, Krause J, Wood AJ. How perceived threat increases synchronization in collectively moving animal groups. Proceedings of the Royal Society B: Biological Sciences. 2010 Oct 22;277(1697):3065–70.

7. Sebanz N, Knoblich G. Prediction in Joint Action: What, When, and Where. Topics in Cognitive Science. 2009;1(2):353–67.

8. Dostálková I, Špinka M. Synchronization of behaviour in pairs: the role of communication and consequences in timing. Animal Behaviour. 2007 Dec 1;74(6):1735–42.

9. Lakin JL, Chartrand TL. Using Nonconscious Behavioral Mimicry to Create Affiliation and Rapport. Psychol Sci. 2003 Jul 1;14(4):334–9.

10. Berscheid E, Walster EH. Interpersonal attraction. Reading, Massachusetts, [etc.: Addison-Wesley; 1977.

11. Hale J, Hamilton AF de C. Cognitive mechanisms for responding to mimicry from others. Neuroscience & Biobehavioral Reviews. 2016 Apr;63:106–23.

12. Oullier O, de Guzman GC, Jantzen KJ, Lagarde J, Scott Kelso JA. Social coordination dynamics: Measuring human bonding. Social Neuroscience. 2008 Jun;3(2):178–92.

13. Hove MJ, Risen JL. It’s all in the timing: Interpersonal synchrony increases affiliation. Social Cognition. 2009;27(6):949–61.

14. Vacharkulksemsuk T, Fredrickson BL. Strangers in sync: Achieving embodied rapport through shared movements. Journal of Experimental Social Psychology. 2012 Jan 1;48(1):399–402.

15. Cornejo C, Hurtado E, Cuadros Z, Torres-Araneda A, Paredes J, Olivares H, et al. Dynamics of Simultaneous and Imitative Bodily Coordination in Trust and Distrust. Front Psychol [Internet]. 2018 [cited 2021 Jan 17];9. Available from: https://www.frontiersin.org/articles/10.3389/fpsyg.2018.01546/full

16. Valdesolo P, Ouyang J, DeSteno D. The rhythm of joint action: Synchrony promotes cooperative ability. Journal of Experimental Social Psychology. 2010 Jul 1;46(4):693–5.

17. Wheatley T, Kang O, Parkinson C, Looser CE. From Mind Perception to Mental Connection: Synchrony as a Mechanism for Social Understanding: Mind Perception and Mental Connection. Social and Personality Psychology Compass. 2012 Aug;6(8):589–606.

18. Cheong JH, Molani Z, Sadhukha S, Chang LJ. Synchronized affect in shared experiences strengthens social connection [Internet]. PsyArXiv; 2020 Jan [cited 2021 Jun 23]. Available from: https://osf.io/bd9wn

19. Marsh KL, Richardson MJ, Schmidt RC. Social connection through joint action and interpersonal coordination. Top Cogn Sci. 2009 Apr;1(2):320–39.

20. Wilson M, Knoblich G. The Case for Motor Involvement in Perceiving Conspecifics. Psychological Bulletin. 2005;131(3):460–73.

21. LaFrance M. Nonverbal Synchrony and Rapport: Analysis by the Cross-Lag Panel Technique. Social Psychology Quarterly. 1979 Mar;42(1):66.

22. Noy L, Dekel E, Alon U. The mirror game as a paradigm for studying the dynamics of two people improvising motion together. PNAS. 2011 Dec 27;108(52):20947–52.

23. Ramseyer F, Tschacher W. Nonverbal synchrony in psychotherapy: Coordinated body movement reflects relationship quality and outcome. Journal of Consulting and Clinical Psychology. 2011;79(3):284–95.

24. Bernieri FJ. Coordinated movement and rapport in teacher-student interactions. J Nonverbal Behav. 1988 Jun 1;12(2):120–38.

25. Konvalinka I, Xygalatas D, Bulbulia J, Schjodt U, Jegindo E-M, Wallot S, et al. Synchronized arousal between performers and related spectators in a fire-walking ritual. Proceedings of the National Academy of Sciences. 2011 May 17;108(20):8514–9.

26. Vesper C, van der Wel RPRD, Knoblich G, Sebanz N. Making oneself predictable: reduced temporal variability facilitates joint action coordination. Exp Brain Res. 2011 Jun;211(3–4):517–30.

27. Konvalinka I, Vuust P, Roepstorff A, Frith CD. Follow you, Follow me: Continuous Mutual Prediction and Adaptation in Joint Tapping. Quarterly Journal of Experimental Psychology. 2010 Nov 1;63(11):2220–30.

28. Eastwood JD, Frischen A, Fenske MJ, Smilek D. The Unengaged Mind: Defining Boredom in Terms of Attention. Perspect Psychol Sci. 2012 Sep 1;7(5):482–95.

29. Zuckerman M. Sensation Seeking and Risk Taking. In: Izard CE, editor. Emotions in Personality and Psychopathology [Internet]. Boston, MA: Springer US; 1979 [cited 2021 Jan 5]. p. 161–97. (Emotions, Personality, and Psychotherapy). Available from: https://doi.org/10.1007/978-1-4613-2892-6_7

30. O’Hanlon JF. Boredom: Practical consequences and a theory. Acta Psychologica. 1981 Oct;49(1):53–82.

31. van Tilburg WAP, Igou ER. On boredom: Lack of challenge and meaning as distinct boredom experiences. Motiv Emot. 2012 Jun 1;36(2):181–94.

32. Hasson U, Frith CD. Mirroring and beyond: coupled dynamics as a generalized framework for modelling social interactions. Philosophical Transactions of the Royal Society B: Biological Sciences. 2016 May 5;371(1693):20150366.

33. Costa VD, Tran VL, Turchi J, Averbeck BB. Dopamine modulates novelty seeking behavior during decision making. Behavioral Neuroscience. 2014;128(5):556–66.

34. Bevins RA, Klebaur JE, Bardo MT. Individual differences in response to novelty, amphetamine-induced activity and drug discrimination in rats. Behav Pharmacol. 1997 Jun;8(2–3):113–23.

35. Wohltjen S, Wheatley T. Eye contact marks the rise and fall of shared attention in conversation. PNAS [Internet]. 2021 Sep 14 [cited 2021 Sep 11];118(37). Available from: https://www.pnas.org/content/118/37/e2106645118

36. Tschacher W, Rees GM, Ramseyer F. Nonverbal synchrony and affect in dyadic interactions. Front Psychol [Internet]. 2014 [cited 2020 Nov 11];5. Available from: https://www.frontiersin.org/articles/10.3389/fpsyg.2014.01323/full

37. Feniger-Schaal R, Hart Y, Lotan N, Koren-Karie N, Noy L. The Body Speaks: Using the Mirror Game to Link Attachment and Non-verbal Behavior. Front Psychol [Internet]. 2018 Aug 23 [cited 2019 Jan 29];9. Available from: https://www.ncbi.nlm.nih.gov/pmc/articles/PMC6115809/

38. Feniger-Schaal R, Schönherr D, Altmann U, Strauss B. Movement Synchrony in the Mirror Game. J Nonverbal Behav. 2021 Mar 1;45(1):107–26.

39. Grammer K, Honda M, Juette A, Schmitt A. Fuzziness of nonverbal courtship communication unblurred by motion energy detection. Journal of Personality and Social Psychology. 1999;77(3):487–508.

40. Shannon CE. A Mathematical Theory of Communication. Bell System Technical Journal. 1948 Oct;27(4):623–56.

41. Varni G, Hupont I, Clavel C, Chetouani M. Computational Study of Primitive Emotional Contagion in Dyadic Interactions. IEEE Transactions on Affective Computing. 2020 Apr;11(2):258–71.

42. Massey FJ. The Kolmogorov-Smirnov Test for Goodness of Fit. Journal of the American Statistical Association. 1951 Mar;46(253):68–78.

43. Granger CWJ. Investigating Causal Relations by Econometric Models and Cross-spectral Methods. Econometrica. 1969 Aug;37(3):424.

44. Keysers C, Gazzola V, Wagenmakers E-J. Using Bayes factor hypothesis testing in neuroscience to establish evidence of absence. Nat Neurosci. 2020 Jul;23(7):788–99.

45. Goss-Sampson M. Bayesian Inference in JASP: a guide for students [Internet]. 2020. 120 p. Available from: http://static.jasp-stats.org/Manuals/Bayesian_Guide_v0_12_2_1.pdf

46. Kass RE, Raftery AE. Bayes Factors. Journal of the American Statistical Association. 1995 Jun;90(430):773–95.

47. van den Bergh D, Clyde MA, Raj A, de Jong T, Gronau QF, Marsman M, et al. A Tutorial on Bayesian Multi-Model Linear Regression with BAS and JASP [Internet]. PsyArXiv; 2020 Apr [cited 2021 Feb 26]. Available from: https://osf.io/pqju6

48. Jiang J, Chen C, Dai B, Shi G, Ding G, Liu L, et al. Leader emergence through interpersonal neural synchronization. Proc Natl Acad Sci USA. 2015 Apr 7;112(14):4274–9.

49. Hart Y, Czerniak E, Karnieli-Miller O, Mayo AE, Ziv A, Biegon A, et al. Automated Video Analysis of Non-verbal Communication in a Medical Setting. Frontiers in Psychology [Internet]. 2016 Aug 23 [cited 2019 Jan 23];7. Available from: http://journal.frontiersin.org/Article/10.3389/fpsyg.2016.01130/abstract

50. Valdesolo P, DeSteno D. Synchrony and the social tuning of compassion. Emotion. 2011;11(2):262–6.

51. Tarr B, Launay J, Dunbar RIM. Music and social bonding: “self-other” merging and neurohormonal mechanisms. Front Psychol [Internet]. 2014 [cited 2021 Feb 18];5. Available from: https://www.frontiersin.org/articles/10.3389/fpsyg.2014.01096/full

52. Nummenmaa L, Glerean E, Viinikainen M, Jaaskelainen IP, Hari R, Sams M. Emotions promote social interaction by synchronizing brain activity across individuals. Proceedings of the National Academy of Sciences. 2012 Jun 12;109(24):9599–604.

53. Burgess AP. On the interpretation of synchronization in EEG hyperscanning studies: a cautionary note. Front Hum Neurosci [Internet]. 2013 [cited 2021 Feb 18];7. Available from: http://journal.frontiersin.org/article/10.3389/fnhum.2013.00881/abstract

54. Rosenblum M, Pikovsky A, Kurths J, Schäfer C, Tass PA. Chapter 9 Phase synchronization: From theory to data analysis. In: Handbook of Biological Physics [Internet]. Elsevier; 2001 [cited 2021 Feb 18]. p. 279–321. Available from: https://linkinghub.elsevier.com/retrieve/pii/S1383812101800129

55. Miller LC. Intimacy and liking: Mutual influence and the role of unique relationships. Journal of Personality and Social Psychology. 1990;59(1):50–60.

56. Sansone C, Thoman DB. Interest as the Missing Motivator in Self-Regulation. European Psychologist. 2005 Jan 1;10(3):175–86.

57. Dahan A, Noy L, Hart Y, Mayo A, Alon U. Exit from Synchrony in Joint Improvised Motion. PLoS One [Internet]. 2016 Oct 6 [cited 2020 Apr 5];11(10). Available from: https://www.ncbi.nlm.nih.gov/pmc/articles/PMC5053605/

58. Fogel A. Developing through relationships: origins of communication, self, and culture. New York: Harvester Wheatsheaf; 1993. 230 p. (The developing body and mind series).

59. Charny JE. Psychosomatic Manifestations of Rapport in Psychotherapy: Psychosomatic Medicine. 1966 Jul;28(4):305–15.

