## Supplementary Material for "Reducing Movement Synchronization to Increase Interest Improves Interpersonal Liking"

**Segmentation.** The segmented mirror games durations were between 84.62 to 127.56 seconds. The time window analysis was performed for a duration of 108.2 seconds, in order to include the entire time series of the vast majority of the games (only four games ended before this point: after 84.62 sec, 100.57 sec, 106.2 sec and 107.74 sec. These dyads are included in the statistics until the mentioned ending point of each). Notice that the last segment is defined as the last movement between two stopping or decelerating points, and hence the partial movement after it is not considered as a movement segment.

**Measuring Synchronization by Maximal Cross-Correlation Instead of Pearson Coefficient Showed Similar Results.** We used multiple linear regression models as well as Bayesian regression models in order to predict dyads' mutual liking by level of synchronization, complexity and novelty. A model including only the level of synchronization predicted 6.1% of variance in liking, $\beta = .25$, $t(98) = 2.53$, $P = .013$, $R^2 = .06$, $BF_{10} = 3.464$ (Supplementary Fig. S1). This result is in line with previous studies (Lakin & Chartrand, 2003), showing the linkage between synchronization and liking. Including also the complexity level significantly improved the model and predicted an additional 6.1% of variance in liking. Accordingly, the linear model with both synchronization and the average entropy as predictors predicted 12.2% of the variance in liking, $F(2, 97) = 6.76$, $P = .002$, $BF_{10} = 17.303$, with a positive correlation between the level of synchronization and liking, $\beta = 0.3$, $t(97) = 3.137$, $P = 0.002$, $\mathrm{BF}_{\mathrm{inclusion}} = 12.976$ and complexity level and liking, $\beta = .254$, $t(97) = 2.6$, $P = .011$, $BF_{inclusion} = 5.924$ (Supplementary Fig. S1b). Introducing novelty to the model did not significantly improve it, and predicted an additional variance of 0.02%, $P = .62$, $BF_{10} = .337$. This result suggests that although the average novelty during the game is related to the synchronization level and the complexity of an interaction, it is not sufficient to create affiliation towards others. Following the comparisons between the models, as demonstrated in Fig. S4b, the best linear model out of the three was:

$$\text{Liking} = -54.79 + 64.4 \times \text{Synchronicity} + 7.77 \times \text{Complexity}$$

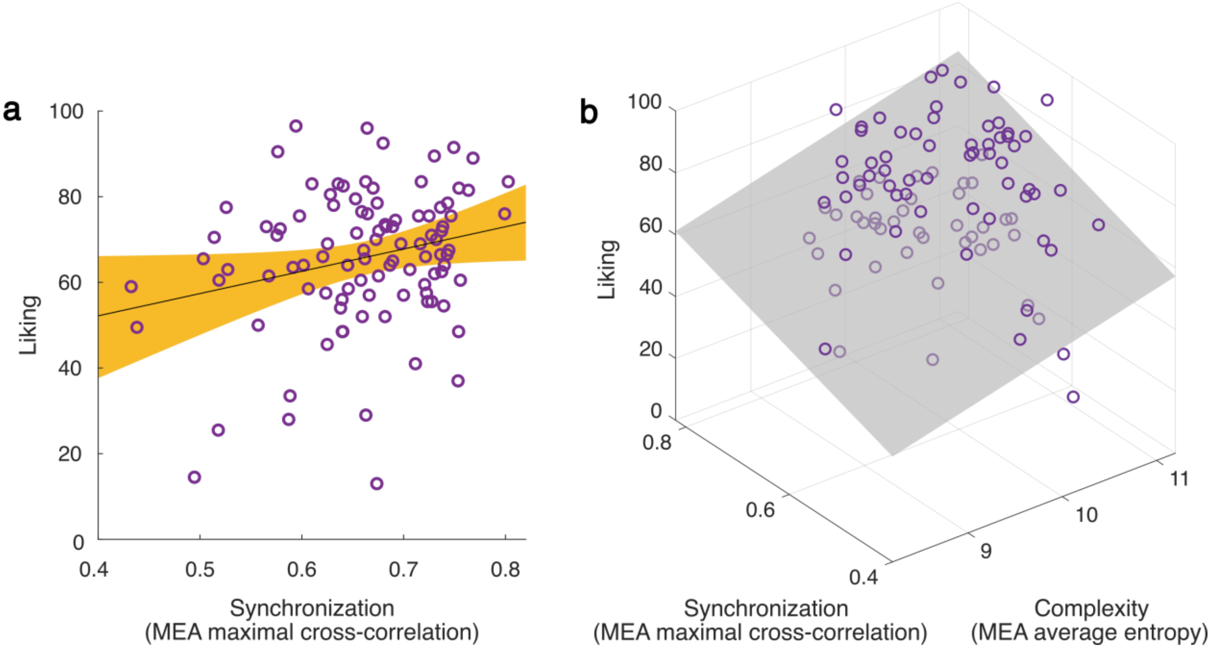


**Supplementary Fig. 1. Predicting liking by dyadic movement features.**

(**a**) Synchronization level of the z-scored MEA signals significantly predict $\beta$ = .25, $t(98)$ = 2.53, $P$ = .013, $R^2$ = .06, $BF_{10}$ = 3.46). Each circle represents a dyad, the black line is the linear regression line, and the orange area marks the confidence interval around the slope of a regression line. (**b**) Multiple linear regression model including both synchronization and complexity significantly improved the model predictions (this model predicted 12.2% of the variance in liking, $R^2$ = .122, $F(2, 97)$ = 6.76, $P$ = .002, $BF_{10}$ = 17.30). Each circle represents a dyad in the 3D space and the grey plane marks the regression surface that was fitted by the model.

Exploring the time domain, synchronization as measured by the maximal cross-correlation showed that in 43.2% out of the total game duration the high-liking group moved in a more synchronized manner than the low-liking group, which is significantly above chance level, binomial $P$ < .001. The high-liking group moved in a significantly more synchronized manner than the low-liking group for 4.96% of the game duration. These results are denoted at the bottom of Supplementary Fig. 2 by the grey and black marks, respectively. In order to show by how much the probability of $h_1$ (the high-liking group is more synchronized than the low-liking group) is expected to be true compared to $h_0$, we performed Bayesian analysis. The dashed line denotes the Bayes factor ($BF_{10}$), which shows substantial Bayesian evidence in 2.5% of the mirror game duration.

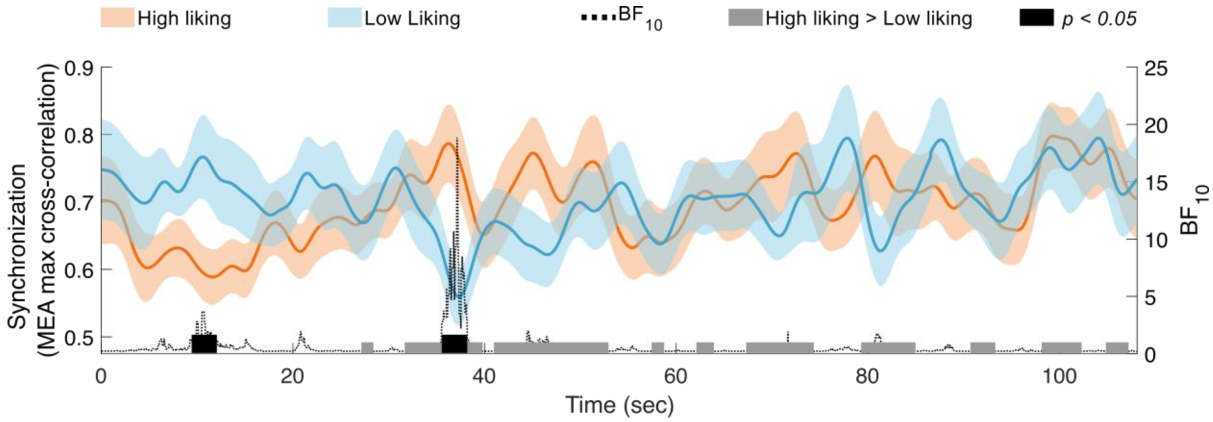


**Supplementary Fig. 2. Maximal cross-correlation along the interaction segments for high and low-liking.** Synchronization was higher in the high-liking group than the low-liking group for 43.2% of the game duration, Binomial sign test $P < .001$. The orange and blue lines denote the average value of the high and low-liking groups respectively, along the mirror game time-points. The shaded orange and blue marks denote the *SE*s of the high and low-liking groups, respectively. The gray marks at the bottom of each panel denote time-points in which the high-liking group had a higher value than the low-liking group. The black marks denote time-points in which there was a significant difference between the liking groups. The dashed lines depict two-tailed Bayes factors ($BF_{10}$).

**Analysis of the role of Synchronization and complexity in mutual liking using stepwise multiple linear regression.** We further conducted a stepwise multiple linear regression in order to examine the optimal contribution of the different movement features (synchronization as measured by Pearson correlation coefficients, synchronization as measured by the maximal cross-correlation, complexity as measured by the average entropy and novelty as was measured by K-S distance) in predicting liking. The final model included the same two predictors as in the original model: synchronization as measured by Pearson correlation coefficients and average entropy. As reported before, this model predicted 14.9% of the variance in liking, $R^2 = .149$, $F(2, 97) = 8.49$, $P < .001$, $BF_{10} = 66.541$, with a positive correlation between the level of synchronization and liking, $\beta = .35$, $t(97) = 3.63$, $P < .001$, and the average entropy and liking, $\beta = .28$, $t(97) = 2.84$, $P = .005$.

**Supplementary Table 1. Summary of the results of the multiple regression models.**

| Model | $R^2$ | *F-Change* (df) | *P*-value | *BF* | Variable | *β* | *t* (df) | *P*-value | *BF* |
|---|---|---|---|---|---|---|---|---|---|
| Synchronization | .08 | 8.28 (1, 98) | | 7.67 | | .28 | 2.88 (1, 98) | .005 | 7.67 |
| Synchronization + Complexity | .15 | 8.49 (2, 97) | < 0.001 | 66.54 | Synchronization | .35 | 3.63 (2, 97) | < .001 | 47.97 |
| | | | | | Complexity | .28 | 2.84 (2, 97) | .005 | 13.86 |
| Synchronization + Complexity + Novelty | .15 | .56 (1, 96) | 0.456 | 24.51 | Synchronization | .37 | 3.7 (1, 96) | < .001 | 34.62 |
| | | | | | Complexity | .25 | 2.42 (1, 96) | .002 | 8.17 |
| | | | | | Novelty | .08 | .75 (1, 96) | 0.456 | 1.03 |
